# Sex-dependent effects of angiotensin type 2 receptor expressing medial prefrontal cortex (mPFC) interneurons in fear extinction learning

**DOI:** 10.1101/2023.11.21.568156

**Authors:** Hannah C. Smith, Zhe Yu, Laxmi Iyer, Paul J. Marvar

**Affiliations:** Department of Neuroscience, George Washington University, Washington, DC; Department of Pharmacology & Physiology, George Washington University, Washington, DC; Department of Psychiatry and Behavioral Sciences, George Washington University, Washington DC

**Keywords:** angiotensin II, PTSD, Fear extinction, AT2R, prefrontal cortex, interneuron

## Abstract

**Background:** The renin-angiotensin system (RAS) has been identified as a potential therapeutic target for PTSD, though its mechanisms are not well understood. Brain angiotensin type 2 receptors (AT2Rs) are a subtype of angiotensin II receptors located in stress and anxiety-related regions, including the medial prefrontal cortex (mPFC), but their function and mechanism in the mPFC remain unexplored. We therefore used a combination of imaging, cre/lox, and behavioral methods to investigate mPFC-AT2R-expressing neuron involvement in fear learning.

**Methods:** To characterize mPFC-AT2R-expressing neurons in the mPFC, AT2R-Cre/td-Tomato male and female mice were used for immunohistochemistry (IHC). mPFC brain sections were stained with glutamatergic or interneuron markers, and density of AT2R^+^ cells and colocalization with each marker was quantified. To assess fear-related behaviors in AT2R-flox mice, we selectively deleted AT2R from mPFC neurons using an AAV-Cre virus. Mice then underwent Pavlovian auditory fear conditioning, approach/avoidance, and locomotion testing.

**Results:** IHC results revealed that AT2R is densely expressed in the mPFC. Furthermore, AT2R is primarily expressed in somatostatin interneurons in females but not males. Following fear conditioning, mPFC-AT2R deletion impaired extinction in female but not male mice. Locomotion was unaltered by mPFC-AT2R deletion in males or females, while AT2R-deleted females had increased exploratory behavior.

**Conclusion:** These results lend support for mPFC-AT2R+ neurons as a novel subgroup of somatostatin interneurons that influence fear extinction in a sex-dependent manner. This furthers underscores the role of mPFC in top-down regulation and a unique role for peptidergic (ie., angiotensin) mPFC regulation of fear and sex differences.

## INTRODUCTION

Post-traumatic stress disorder (PTSD) is a strong predictor of cardiovascular disease (CVD), the primary cause of death in both men and women in the United States. Nevertheless, the specific biological, behavioral, and causal mechanisms that connect PTSD with this co-occurrence of CVD, particularly with notable differences between genders, remain unclear(1–4). Given its involvement in maintaining cardiovascular balance and responding to stress-related emotions, the renin-angiotensin system (RAS) has emerged as a potentially significant bridge between these conditions(5–9).

Angiotensin II serves as the main functional molecule of the RAS, and its effects are facilitated by binding to its primary receptor subtypes, the angiotensin type 1 (AT1R) and type 2 (AT2R) receptors(10). These receptors are expressed throughout the brain, with more extensive research focused on their presence in regions like the hypothalamus, brainstem, and forebrain. However, AT1R and AT2R are also expressed in cortical-limbic brain structures, such as the amygdala and medial prefrontal cortex (mPFC), which are crucial regions involved in fear memory and learning(11–15). Previous research studies, including ours, have highlighted the unique expression of neuronal AT2R in the amygdala and prefrontal cortex and their distinct roles in threat-related memory(11,12). Their functional mechanism in the mPFC, however, remains largely unknown.

The role of the brain AT2R is increasingly being studied in cardiovascular(13,16,17) as well as emotional(11,18–20) regulation. As a G_i_-protein coupled receptor, AT2R has been shown in multiple brain regions to decrease the firing rate and neuronal activity of both excitatory and inhibitory neurons(21). This inhibitory effect influences physiological outcomes of stress, acting both neuroprotective and hypotensive(14,22), as well as behavioral outcomes of stress, as activation of central amygdala AT2R decreases fear expression while whole-brain knockout increases anxiety in mice (11,19). Interestingly, while most studies examine these effects in male mice, AT2R is an X-linked gene upregulated by estrogen signaling(10,23,24), and some studies have shown that AT2R knockout causes cognitive deficits in only female mice(25), indicating a potentially sex-specific function. Though AT2R is highly expressed in the mPFC(12,15), a region integral to both behavioral and physiological outcomes of fear, no studies to date have examined what cells these receptors are located on, their behavioral function, or their mechanism of action.

The mPFC is a robust top-down regulator of learning and memory, and acts via a complex network of microcircuits wherein different classes of interneurons regulate the firing of excitatory projection neurons to downstream brain regions such as the amygdala, brainstem, and thalamus(26). These regulatory projections are hypoactive in PTSD, and reestablishing their activity can rescue impaired extinction and disordered fear learning(27–32). Clinical studies have implicated that the RAS plays an important role in this top-down fear regulation by the mPFC; treatment with AT1R antagonist losartan in humans facilitated fear extinction, while concurrent fMRI showed that losartan increased mPFC connectivity to the basolateral amygdala (BLA)(33). Due to the opposing actions of AT1R and AT2R, this result of AT1R antagonism indicates a potential role for mPFC AT2R in the top-down regulation of fear extinction in clinical as well as preclinical studies.

In this study, we aimed to characterize the expression and functional role of AT2R in the mPFC. In doing so, we provide a novel understanding of AT2R’s mechanism and the cells it acts on, as well as its downstream behavioral targets in the regulation of fear learning and memory. By using male and female mice we investigate the how AT2R’s female-biased effects influence both neurobiology and fear-related behavior; this factor is especially important given that women are twice as likely to develop PSTD as men(4,34,35). By combining immunohistochemistry, cre/lox techniques, and classic Pavlovian fear conditioning, we identify a novel population of AT2R-expressing interneurons in the mPFC that influence fear learning and memory in a sex-dependent manner, establishing a potential therapeutic target for disordered fear learning in females.

## METHODS AND MATERIALS

**Animals:** All experimental procedures were approved by the Institutional Care and Use Committee (IACUC) of the George Washington University and followed National Institutes of Health guidelines. 8–10-week-old male and female transgenic mice were used for the following studies and were housed in a temperature and humidity-controlled room on a 12hr light/dark cycle. Food and water were available ad libitum. All mouse lines are on a C57/BL/6 background and are summarized in **Table 1**.

**Table 1.** Experimental mouse models used. A summary of transgenic mouse models used in this study, their genetic backgrounds, and where they were received from.

| Abbreviation | Genetic Information | Received From |
| --- | --- | --- |
| AT2R-flox | *Agtr2*<sup>loxP/y</sup> | University of Florida |
| AT2R-cre | B6.Cg-*Agtr2*<sup>em1(cre)Adk</sup>/J | University of Florida |
| Ai14 | B6;129S6-*Gt(ROSA)26Sor*<sup>tm14(CAG-tdTomato)Hze</sup>/J | Jackson Laboratories 007908 |
| AT2R-cre/td-Tomato | Breeding AT2R-cre with Ai14 |  |

**Immunohistochemistry:** Immunohistochemistry was used to validate injection location of AT2R-flox mice as well as to characterize AT2R-expressing mPFC neurons in AT2R-cre/td-Tomato reporter mice. Mice were anesthetized with urethane (275 mg/ml, Thermo Fisher Scientific, MA) and perfused with 4% paraformaldehyde (Electron Microscope Science, PA). Brains were removed and post-fixed overnight in the same solution, then transferred to 30% sucrose for 2 days for dehydration. Brains were then embedded in optimal cutting temperature (OCT) compound (Thermo Fisher Scientific, MA) and stored overnight in -80°C. A cryostat (CryoStar NX 50, Thermo Fisher Scientific, MA) was used to section brains into 30μM free-floating serial brain sections. Sections were washed in phosphate buffer saline (PBS) for 15 min and blocked with 5% normal donkey serum, 5% bovine serum albumin, and 0.3% Triton-X-100 in PBS for 1h at room temperature. Primary antibodies (**Table 2**) were added to the solution and brain sections incubated for 24h at 4°C. Sections were then rinsed (3×15min) in PBS and incubated in corresponding secondary antibodies (Thermo Fisher Scientific, MA) for 2h at room temperature. Following a final series of rinses (3×15min), sections were mounted on Superfrost Plus slides (Thermo Fisher Scientific, MA) and air dried before being cover-slopped with ProLong® Diamond Antifade Mountant (Thermo Fisher Scientific, MA). After staining, sections were imaged using 25x water immersion objective on a Zeiss Spinning Disk Confocal microscope. Colocalization was quantified by a blinded researcher using Zeiss Microscope Software ZEN 2 (Carl Zeiss, Germany). AT2R/tdTomato+ cell bodies were identified, counted, and marked, and then colocalization with each marker was assessed by switching microscope channels and identifying and counting each stained cell body and whether it was also marked as AT2R+. **Animal Surgery:** Ketamine (82.5 mg/kg) and xylazine (12.5 mg/kg) anesthesia were intraperitoneally injected. Cre or GFP expressing adeno-associated virus (GFP pAAV.CMV.PI.EGFP.WPRE.bGH, Addgene 105530; Cre pENN.AAV.CMVs.PI.Cre.rBG, Addgene 105537) were bilaterally injected into the mPFC of AT2R-flox mice at 2.5mm caudal, ±0.3mm lateral to bregma, and 2.1mm below the skull surface with an UltraMicroPump III and microprocessor controller (World Precision Instruments, FL). 400nL was injected bilaterally at a rate of 100nL/min. Following surgery, the mice were group housed in their home cages for a 3-week postoperative period.

**Table 2.** Antibodies used for immunohistochemistry. A summary of antibodies used for immunohistochemistry co-staining analysis, as well as the manufacturer, species, and dilution for each.

| **Antibody** | **Manufacturer (catalog #)** | **Species** | **Dilution** |
| --- | --- | --- | --- |
| Anti-GFP | Abcam (ab13970) | Goat | 1:2000 |
| Anti-TBR1 | Abcam (ab31940) | Rabbit | 1:500 |
| Anti-Calretinin | Abcam (ab702) | Rabbit | 1:100 |
| Anti-Calbindin | Abcam (ab75524) | Mouse | 1:1000 |
| Anti-nNos | Immunostar (24287) | Rabbit | 1:2000 |
| Anti-Parvalbumin | Abcam (ab11427) | Rabbit | 1:2000 |
| Anti-Somatostatin | Immunostar (20067) | Rabbit | 1:2000 |

**mRNA extraction and RT-qPCR:** Mice were sacrificed and brains were collected and flash frozen. The mPFC was collected using a 1mm diameter brain tissue punch. Total RNA was extracted from the punches using Trizol reagent (Thermo Fisher, Waltham, MA) according to the manufacturer’s instructions and one microgram of RNA was reverse transcribed to cDNA using qScript cDNA Supermix (QuantBio, Beverly, MA). Gene expression changes for 18S and AT2R were detected using Taqman primers (18S (Mm04277571_s1), AT2R (Mm1341373_m1), Thermo Fisher) and the Applied Biosystems^TM^ ViiA Real-Time PCR Systems by relative quantitative RT-PCR (Thermo Fisher Scientific, Waltham, MA; Applied Biosystems, Foster City, CA) and expression fold change was calculated using the ΔΔCq calculation method, normalizing to loading housekeeping gene 18S control.

**Cue-dependent Pavlovian fear conditioning and extinction:** Pavlovian fear conditioning, as previously described by our lab(7,9,11), was used to determine the effects of mPFC AT2R deletion on conditioned fear behavior by pairing an auditory cue with a light foot shock. Mice were habituated to the fear conditioning chamber for two days (20min, 40min) before undergoing aversive associative conditioning on experimental day 3, during which mice received 5 conditioned stimulus (CS)/unconditioned stimulus (US) pairings using a 30s auditory cue (6kHz, 75dB, CS) co-terminating with a mild foot shock (0.5s, 0.6mA, US) at an inter-trial interval of 5min. 24hrs later (experimental day 4), mice were placed in a novel context and underwent extinction training consisting of a 5min pre-CS period followed by 30 CS presentations (30s each, 30s inter-trial interval). The same protocol was repeated on experimental day 5 in the same context as extinction training to test retention of fear extinction. 7 days after fear conditioning (experimental day 10), mice were again placed in the novel context and the CS was presented 40 times. The mice were recorded and freezing calculated using FreezeFrame 3.0 software (Actimetrics, Wilmette, IL).

**Generalized anxiety measures:** Open field (OF) and elevated plus maze (EPM) tests were used as described previously by our lab(11) to determine whether deletion of mPFC AT2Rs affected locomotion or anxiety-like behavior. These tests were performed one week after fear conditioning and extinction testing. In the OF test, mice were placed in the center of the open field arena (35cm x 35cm x 35cm, opaque plexiglass) and allowed to freely explore for 30min. Locomotion, speed, and position in the apparatus were recorded and analyzed using Anymaze software (ANY-Maze, Wood Dale, IL). In the EPM test, mice were placed in the center of the apparatus facing the same closed arm and allowed to freely explore for 5min. Total arm entries, open arm entries, and percentage of time spent in the open arms were analyzed using Anymaze software (ANY-Maze, Wood Dale, IL).

**Data presentation and statistical analysis:** All data analysis was completed using GraphPad Prism 9.0 software (GraphPad Software, CA) and checked for outliers using the ROUT outlier test (Q=1%). In all data reported, a *p* value of *p*<0.05 is considered statistically significant. Mean differences between groups were compared using unpaired t-tests and two-way RM ANOVA with Holm-Šídák post-hoc analyses. To assure rigor, reproducibility, and minimal use of animals, power analyses were conducted to determine a sample size providing 80% power and significance at a level of 5%.

## RESULTS

**AT2R-cre/td-Tomato^+^ cells are densely expressed throughout the mPFC of males and females:** Using an AT2R-Cre mouse cross-bred with a td-Tomato reporter mouse (**Fig. 1A**) we examined AT2R-tdTomato+ expression in the mPFC of male and female mice using immunohistochemistry (IHC). Both males and females had equally dense expression of AT2R in the PL and IL of the mPFC (**Fig. 1B**; males: 208.6 ± 48.8 cells/mm^2^, females: 139.0 ± 59.8 cells/mm^2^, p=0.42) throughout cortical layers L2/3-L6 (**Fig. 1C**). This density did not change across rostral-caudal depth of the mPFC (**Fig. 1D**) in males or females (**Fig. 1E**). There was little AT2R expression observed in L1 in males or females throughout the mPFC (**Fig. 1C,F**). This dense and nonspecific expression of AT2R^+^ cells through the mPFC indicates that they may contribute to mPFC top-down regulation of fear learning(27,29–31).

**Figure 1.**
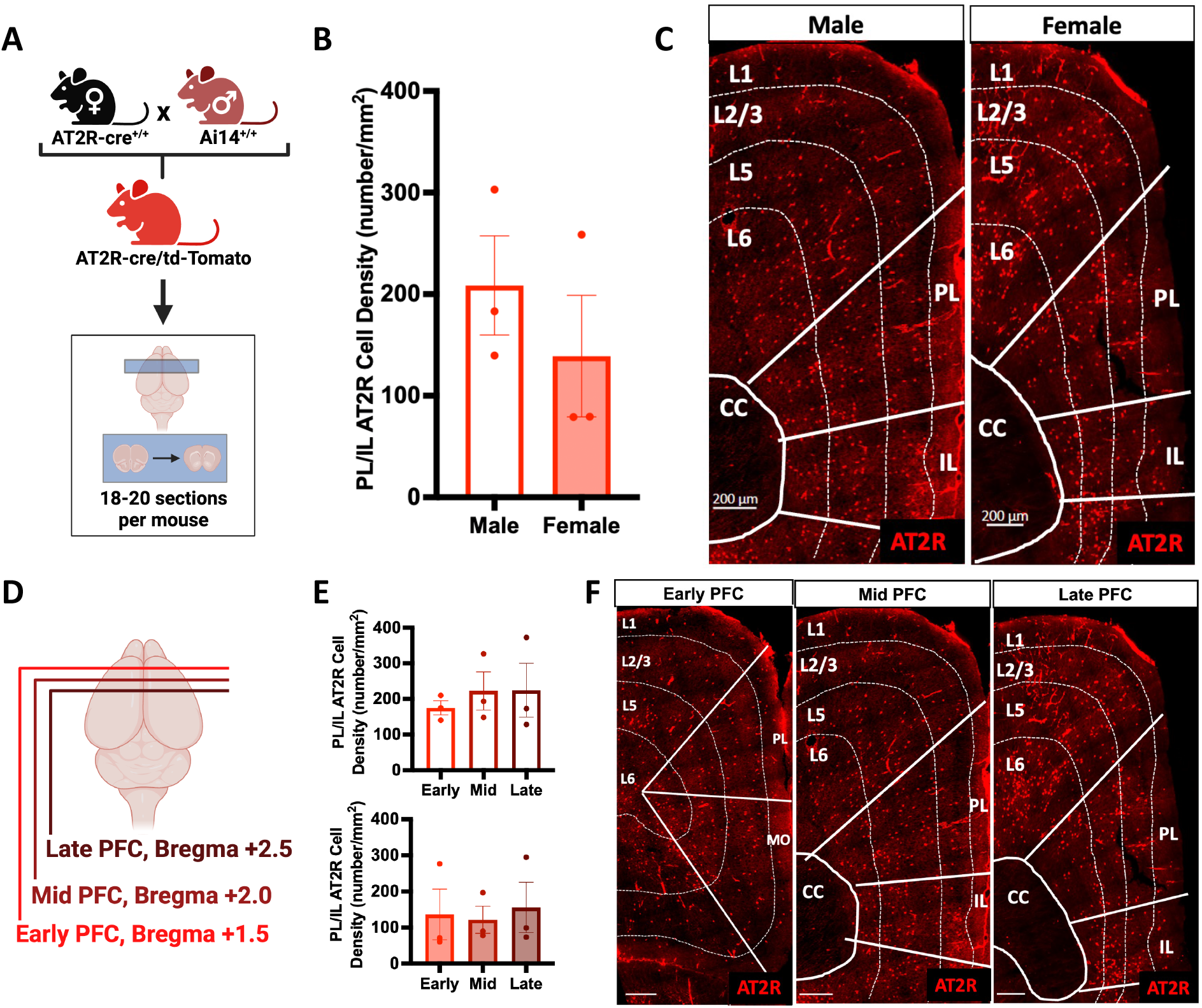
AT2R/td-Tomato^+^ mPFC expression in male and female mice. **A.** Generation of the AT2R-cre/td-Tomato reporter mouse and experimental approach to test mPFC AT2R distribution with immunohistochemistry (IHC). **B**. AT2R+ cells are equally expressed in the prelimbic (PL) and infralimbic (IL) regions of the mPFC of male (n=3) and female (n=3) mice (t(4)=0.90, p=0.42). **C.** Representative coronal sections showing laminar distribution of AT2R^+^ neurons in layers 2/3, 5, and 6 of the mPFC of male and female mice. **D**. Approach for assessing distribution of AT2R^+^ neurons through the depth of the mPFC. **E**. Rostral/caudal depth of mPFC had no impact on distribution of AT2R^+^ neurons in male (early mPFC, 175.0 ± 20.0 cells/mm^2^, mid mPFC, 222.6 ± 53.7 cells/mm^2^, late mPFC, 224.4 ± 75.2 cells/mm^2,^ F(2,6)=0.26, p=0.78) or female (early mPFC, 136.2 ± 70.4 cells/mm^2^, mid mPFC, 121.4 ± 37.7 cells/mm^2^, late mPFC, 155.7 ± 70.0 cells/mm^2^ F(2,6)=0.08, p=0.93) brains. **F**. Representative coronal sections showing laminar distribution of AT2R^+^ neurons throughout rostral/caudal mPFC. All scale bars indicate 200µm.

**Distribution of AT2R-expressing neurons across mPFC cell types:** Though the mPFC is primarily populated by glutamatergic projection neurons(26), AT2R-tdTomato+ co-staining with glutamatergic marker TBR1 (**Fig. 2A**) revealed sparse colocalization in the PL and IL of both males and females (**Fig. 2B-E**; males: 7.1% ± 1.8%; females: 13.2% ± 4.2%, p=0.25). This low percentage of co-staining indicates that AT2R in the mPFC is likely acting via a microcircuit(29,36–38), rather than directly on mPFC projection neurons.

**Figure 2.**
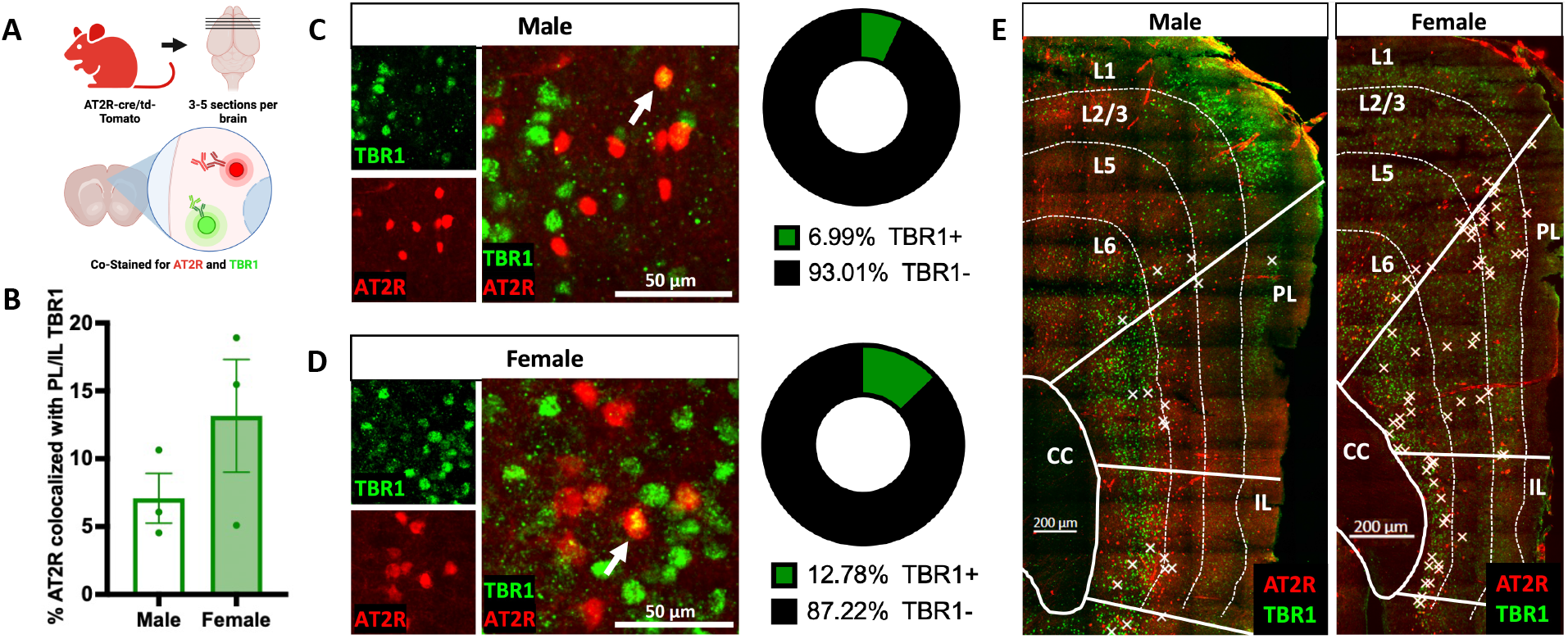
Low expression of AT2R/td-Tomato^+^ on excitatory mPFC neurons. **A**. Experimental approach for co-staining of AT2R-tdTomato^+^ cells with TBR1 neurons in the mPFC (n=3/group). **B**. Sex does not significantly affect AT2R colocalization with TBR1 in the PL and IL of the mPFC (t(4)=1.34, p=0.25). **C-D.** Left, representative coronal section from the male (**C**) and female (**D**) mPFC. AT2R-tdTomato and TBR1 signals are colocalized (indicated with arrows). Right, Percentage of AT2R^+^ neurons colocalized with TBR1 in the mPFC of male (**C**) and female (**D**) mice. **E.** Representative coronal sections from male (left) and female (right) mPFC showing laminar distribution of AT2R/TBR1 colocalizations (indicated with X).

To further characterize the cells on which AT2R is expressed in the PL and IL of the mPFC, IHC was used for co-staining analysis of mPFC sections taken from male and female AT2R-cre/tdTomato mice (**Fig. 3A**, n=3/group). Colocalization was examined between AT2R-tdTomato^+^ cells and 6 cell type markers: somatostatin (som) interneurons, parvalbumin (PV) interneurons, neuronal nitric oxide synthase (nNos) interneurons, calbindin (CB) interneurons, calretinin (CR) interneurons, and TBR1 glutamatergic projection neurons. Sex did not impact AT2R^+^ cell distribution on PV interneurons (**Fig. 3B**), nNos interneurons (**Fig. 3C**), CB interneurons (**Fig. 3D**), or CR interneurons (**Fig. 3E**). Across males and females, there is relatively minimal colocalization through the mPFC of AT2R-tdTomato^+^ cells with these four interneuron markers (**Fig. 3B-E**; PV 5.60%, nNos 4.30%, CB 18.30%, CR 6.20%).

**Figure 3.**
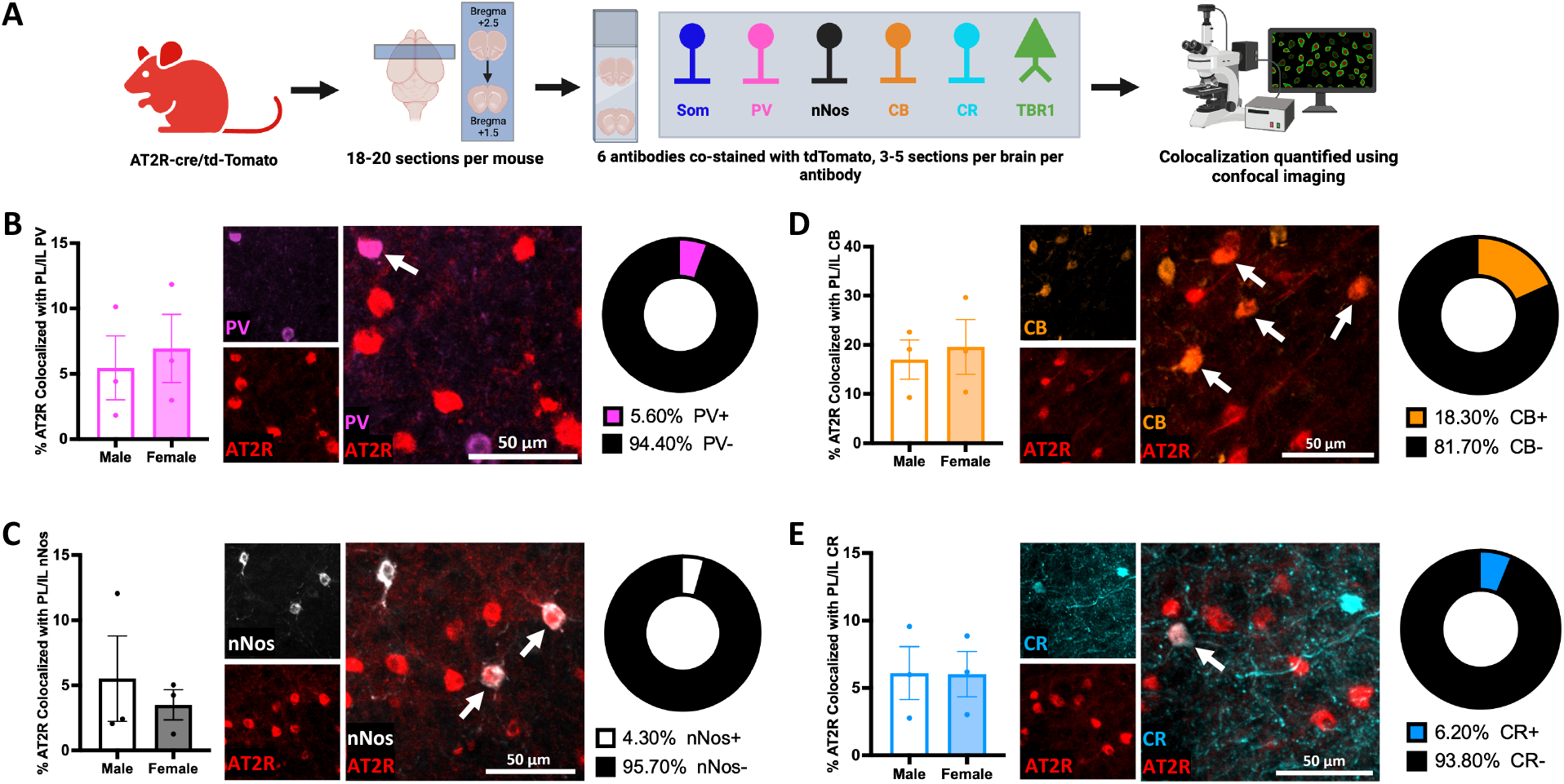
Characterization of AT2R/td-Tomato^+^ in mPFC cell types. **A**. Experimental approach to test mPFC-AT2R^+^ cell type in the mPFC of male and female mice (n=3/group). **B-E**. Sex has no effect on AT2R colocalization with PV interneurons (**B**, male: 5.5% ± 2.5% colocalized, female: 6.9% ± 2.6% colocalized, t(4)=0.41, p=0.70), nNos interneurons (**C,** male: 5.5% ± 3.3% colocalized, female: 3.5% ± 1.2% colocalized, t(4)=0.58, p=0.59)), CB interneurons (**D**, male: 17.0% ± 4.0% colocalized, female: 19.6% ± 5.6% colocalized, t(4)=0.38, p=0.72), or CR interneurons (**E**, male: 6.1% ± 1.9% colocalized, female: 6.0% ± 1.7% colocalized, t(4)=0.03, p=0.98). PL, prelimbic cortex of the mPFC; IL, infralimbic cortex of the mPFC; Som, somatostatin; PV, parvalbumin; nNos, neuronal nitric oxide synthase; CB, calbindin; CR, calretinin. Colocalization indicated with arrows.

**mPFC AT2R is primarily found on somatostatin interneurons in females:** Colocalization analysis between AT2R and somatostatin in mPFC brain sections from male and female AT2R-cre/tdTomato mice (**Fig. 4A**) revealed that somatostatin is the primary AT2R-expressing cell type in females and males, and is more densely colocalized in the PL/IL of females than males (**Fig. 4B-D**; males: 16.5% ± 1.2%, females: 31.8% ± 4.2%, p=0.02). In the PL of both males and females, somatostatin colocalization is distributed throughout L2/3-L6, while in the IL colocalization is biased towards L5 and L6 (**Fig. 4E**). Because somatostatin interneurons in the mPFC have a known regulatory role in fear memory and extinction(39–42) and are activated by stress in a sex-dependent manner(43), we sought to examine the functional role of these AT2R-expressing neurons in fear extinction of male and female AT2R-flox mice.

**Figure 4.**
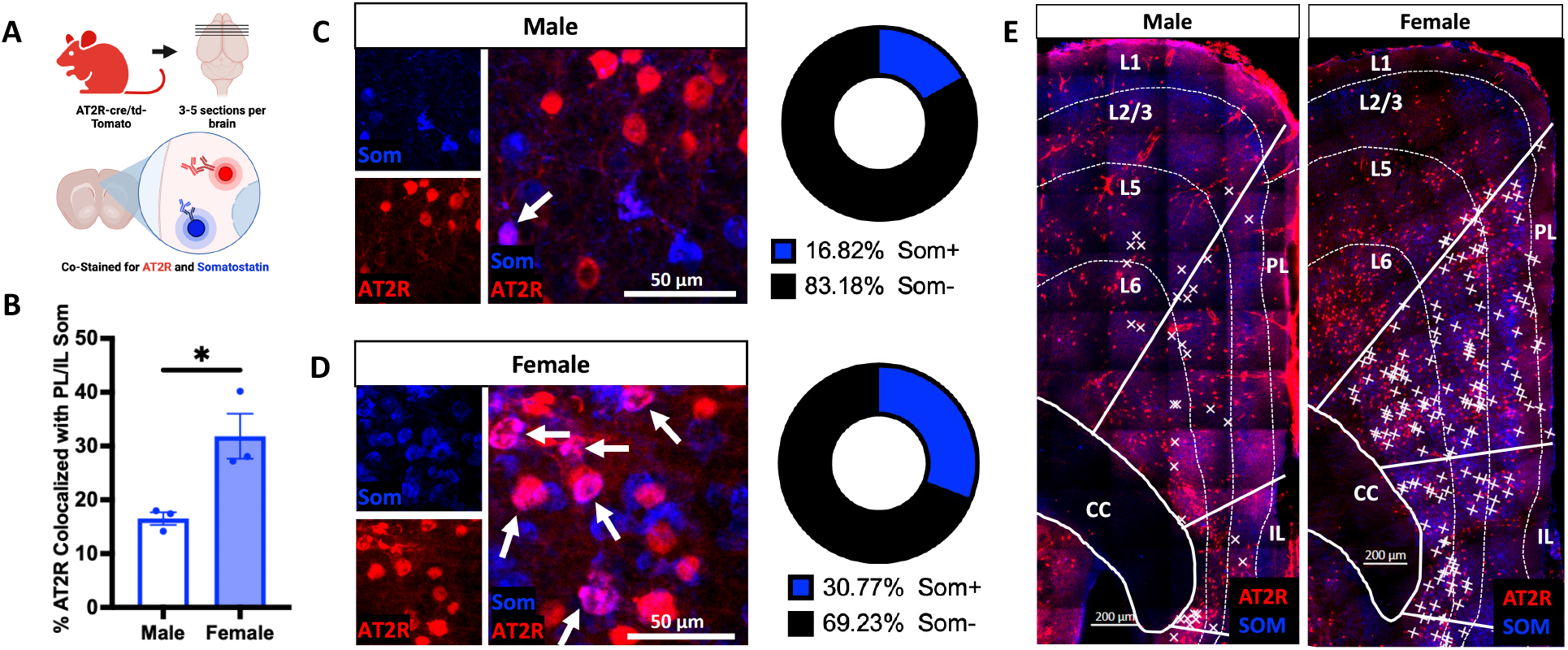
High expression of AT2R/td-Tomato^+^ on mPFC somatostatin interneurons in female mice. **A**. Experimental approach for co-staining of AT2R-tdTomato^+^ cells with somatostatin neurons in the mPFC (n=3/group). **B**. Females have significantly greater AT2R/somatostatin colocalizations in the PL and IL of the mPFC (*t(4)=3.52, p=0.02*). **C-D.** *Left*, representative coronal section from the male (**C**) and female (**D**) mPFC. AT2R-tdTomato and somatostatin signals are colocalized (indicated with arrows). *Right*, Percentage of AT2R+ neurons colocalized with somatostatin in the mPFC of male (**C**) and female (**D**) mice. **E.** Representative coronal sections from male (*left*) and female (*right*) mPFC showing laminar distribution of AT2R/somatostatin colocalizations (indicated with X).

**mPFC AT2R deletion inhibits fear extinction learning in females, but not males:** To assess the role of mPFC AT2R in fear learning, AT2R^flox/flox^ male and female mice received either Cre-expressing adeno-associated virus (AAV-Cre) or GFP-expressing adeno-associated virus (AAV-GFP) into the mPFC to selectively delete AT2R. Four weeks post-injection, AT2R gene expression was significantly decreased in AAV-Cre injected mice as compared to the AAV-GFP control group, confirming site-specific AT2R deletion in the mPFC (**Fig. 5B-C**, *Males:* p=0.02; *Females:* p<0.01). Once mPFC-AT2R deletion was confirmed, we used Pavlovian fear conditioning (5 CS-US pairings) and extinction (3 days, 30 CS per day) to determine the impact on fear behavior (**Fig. 5A**). In males, AT2R deletion had no impact on fear conditioning or freezing to the CS during extinction testing (p=0.76, **Fig. 5D**).

**Figure 5.**
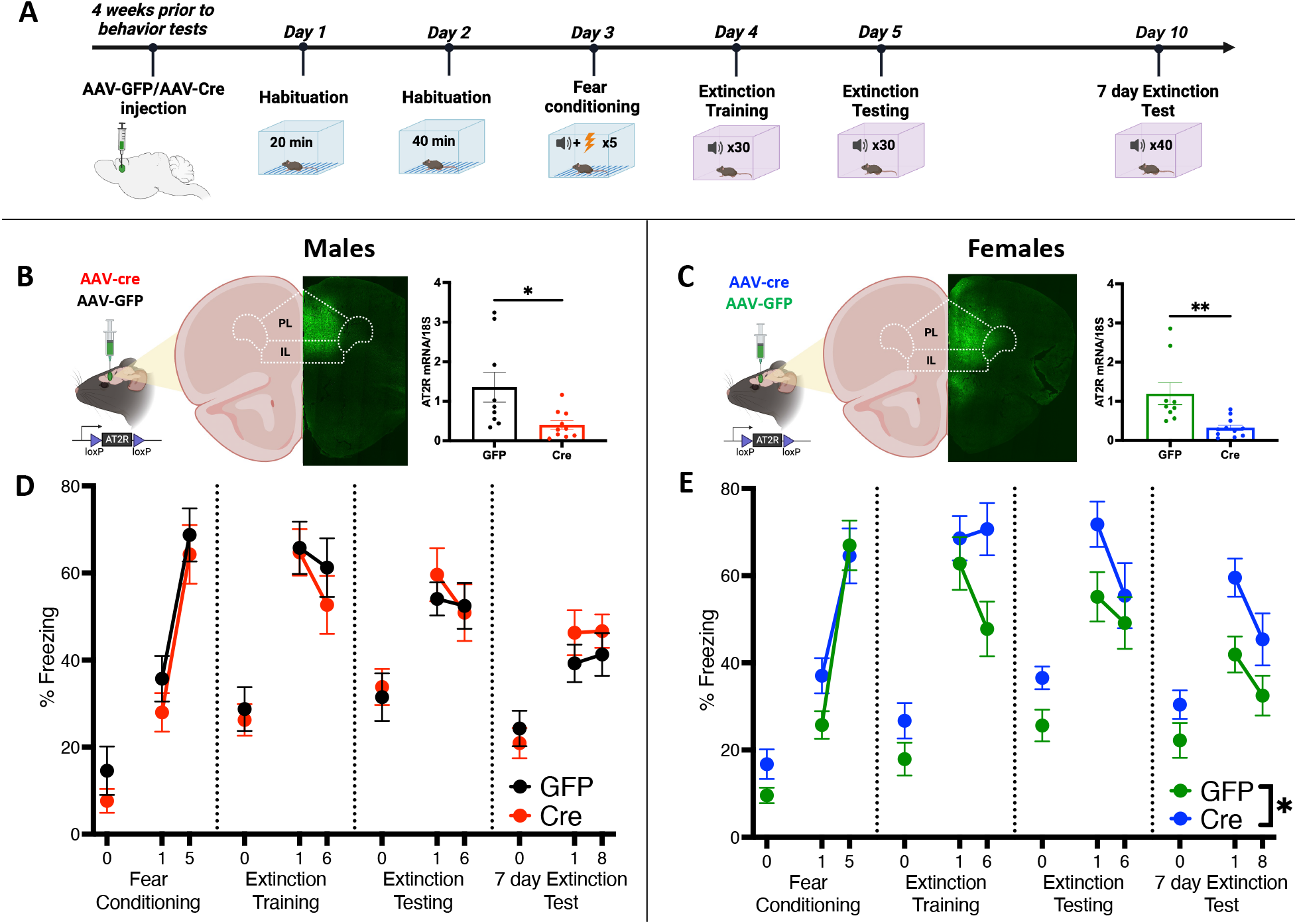
AT2R cre/lox mPFC deletion delays extinction learning in females, but not males. **A.** Experimental protocol for the auditory cue-dependent fear extinction test. **B.** *Left*, strategy for injection of AAV-cre or AAV-GFP into the mPFC of male AT2R-flox mice and representative injection image in the mPFC. *Right*, injection of AAV-cre into the mPFC successfully decreased expression of AT2R in males (GFP 1.35 ± 0.38 fold change, Cre 0.40 ± 0.11 fold change, t(17)=2.54, p=0.02). **C.** *Left*, strategy for injection of AAV-cre or AAV-GFP into the mPFC of female AT2R-flox mice and representative injection image in the mPFC. *Right*, injection of AAV-cre into the mPFC successfully decreased expression of AT2R in females (GFP 1.20 ± 0.28 fold change, Cre 0.32 ± 0.07 fold change, t(18)=3.28, p<0.01). **D.** AT2R deletion from the male mPFC does not impact fear acquisition or extinction learning across three testing days in males (F(1,17)=0.09, p=0.76) **E.** In females, AT2R deletion from the mPFC significantly inhibits extinction learning (F(1,20)=5.35, p=0.03). n=9-12/group.

In females, however, AT2R deletion significantly inhibited extinction learning across three days (p=0.03, extinction training, extinction testing, and a 7-day test of long-term extinction retention). Cre-injected females displayed higher freezing across all days of extinction testing. Together, these results show that AT2R deletion from the mPFC significantly delays both short- and long-term extinction learning in female, but not male, AT2R-flox mice.

**Effects of mPFC AT2R deletion on locomotion, approach/avoidance behavior, and exploration:** To determine whether the difference in fear extinction observed with AT2R-mPFC deletion may be a result of off-target effects on anxiety, locomotion, or a deficit in exploratory behavior, male and female GFP- and Cre-injected AT2R-flox mice underwent open field (**Fig. 6A,E**; OF) and elevated plus maze(**Fig. 6I,M**; EPM) tests. mPFC AT2R deletion did not affect total distance traveled in the OF in males (**Fig. 6B**; GFP 44.11m ± 5.92m, Cre 56.00m ± 10.29m, p=0.35) or females (**Fig. 6F**; GFP 42.86m ± 4.85m, Cre 42.39m ± 4.97m, p=0.95), or total arm entries in the EPM in males (**Fig. 6J**; GFP 26.75 ± 1.45, Cre 31.67 ± 2.17, p=0.09) or females (**Fig. 6N**; GFP 31.20 ± 2.18, Cre 36.22 ± 3.96, p=0.27). While mPFC AT2R deletion did not impact anxiety-like measures in the OF in males (**Fig. 6C-D**; center entries: GFP 70.75 ± 13.99, Cre 106.6 ± 18.87, p=0.16; time in center: GFP 117.60s ± 27.33s, Cre 175.20s ± 27.85s, p=0.16) or females (**Fig. 6G-H**; center entries: GFP 62.60 ± 8.40, Cre 70.44 ± 11.06, p=0.58; time in center: GFP 88.38s ± 13.22s, Cre 113.00s ± 18.67s, p=0.29) or approach, avoidance or exploratory behavior in the EPM in males (**Fig. 6K-L**; open arm entries: GFP 12.38 ± 1.02, Cre 13.22 ± 1.54, p=0.66; time in open arm: GFP 43.25s ± 5.88s, Cre 40.73s ± 6.27s, p=0.78), AT2R deletion from the mPFC increased some measures of exploratory behavior in the EPM in females (**Fig. 6K-L**; open arm entries: GFP 12.50 ± 0.96, Cre 20.00 ± 2.81, p=0.02; time in open arm: GFP 44.54s ± 8.37s, Cre 64.99s ± 12.62s, p=0.19). Taken together, these results show that AT2R deletion from the mPFC does not affect locomotion in males or females but may increase some aspects of exploratory behavior in females.

**Figure 6.**
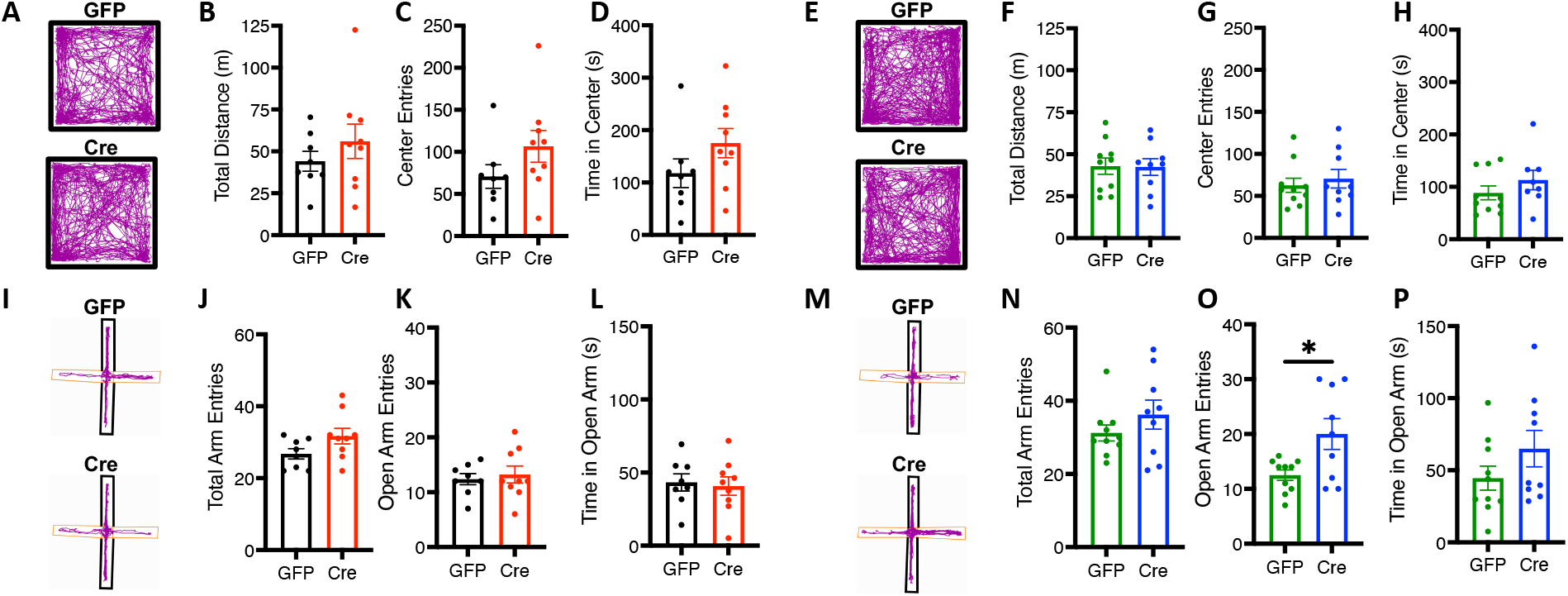
Effects of AT2R cre/lox mPFC deletion on behavioral assays of locomotion and exploration in males and females. **A,E.** Representative locomotor traces of GFP- and Cre-injected males and females in the open field (OF) test. **A-H.** AT2R deletion has no impact on locomotion (**B**, t(15)=0.97, p=0.35 **F** t(17)=0.07, p=0.95, total distance traveled) or generalized anxiety (**C** t(15)=1.49, p=0.16, **G** t(17)=0.57, p=0.58, center entries; **D**, t(15)=1.47, p=0.16, **H** t(17)=1.11, p=0.29, time in center) in males or females. **I,M**. Representative locomotor traces of GFP- and Cre-injected males and females in the elevated plus maze (EPM) test. **I-P**. AT2R deletion has no impact on exploratory behavior (**J**, total arm entries t(15)=1.84, p=0.09) or generalized anxiety in males (**K**, open arm entries t(15)=0.45, p=0.66; **L** t(15)=0.29, p=0.78, time in open arm). mPFC AT2R deletion increases some aspects of exploratory behavior in females (**N**, total arm entries, t(17)=1.14, p=0.27; **O**, open arm entries, t(17)=2.64, p=0.02; **P**, open arm time, t(17)=1.38, p=0.19. OF, open field; EPM, elevated plus maze. n=8-10/group.

## DISCUSSION

Significant ongoing research on post-traumatic stress disorder (PTSD) is directed towards enhancing translational preclinical models with the aim of developing therapies that are efficacious in humans (44,45). This evolution of preclinical models is especially important when investigating sex differences in PTSD, given the lack of therapies developed using female subjects(35,46). Many studies are focused on the neuronal circuitry of PTSD in therapy development, as pharmacologically targeting the conditioned threat memory circuits involved in fear extinction, for example, may benefit exposure-based therapies, one of the most widely used treatment strategies for PTSD(28,29,33,47–49). Further understanding sex-based differences in neuronal circuits may also provide insight into developing therapies effective in both males and females(34,35,46,50). In line with this, recent preclinical and clinical studies have identified the brain RAS as a potential drug target for extinction-based therapies, and have begun investigating the neuronal mechanisms it may be working through(7,8,11,33,51).

Previous studies in mice have shown that manipulation of brain AT2R can influence cognition and fear memory as well as physiological and cardiovascular stress, potentially in a sex-dependent manner(11,13,20,23,52–55). Pharmacological activation of AT2R in the central amygdala decreases fear expression(11), while knockout of AT2R results in anxiety-like behavior and cognitive deficits, particularly in females(52–54). In the hypothalamus, activation of AT2R attenuates hypertension by decreasing the firing of pressor neurons, a protective effect that, similar to the behavioral findings, is also more prominent in females(23,55).

Studies involving AT1R can also inform the functional role of AT2R, as AT1R and AT2R serve opposing physiological roles. Preclinical studies from our lab and others have shown that systemic AT1R blockade with losartan as well as central-amygdala-specific AT1R deletion enhance extinction learning(7,56,57). In clinical studies, losartan improves extinction, decreases PTSD symptom severity, and increases the top-down control of the mPFC over the amygdala(33,58). As evidence shows that losartan upregulates AT2R, this indicates that the benefit of AT1R blockade in both humans and mice may be via an AT2R-dependent mechanism(59,60).

While immunostaining studies have shown that AT2R is highly expressed in the mPFC(12,15), no research to date has investigated the function of mPFC AT2R or how its activity affects top-down regulation of fear learning. Here, we identify a novel population of somatostatin AT2R-expressing interneurons in the mPFC and show the contribution of AT2R^+^ mPFC neurons to the extinction of learned fear in females, proposing a potential therapeutic target for delayed fear extinction in PTSD.

To first classify the cell types expressing AT2R in the mPFC, we crossed AT2R-cre mice(55) with Ai9 tdTomato reporter mice. Consistent with previous immunostaining studies(12,15), we found high density of AT2R-expressing cells throughout the prelimbic (PL) and infralimbic (IL) cortices of the mPFC (**Fig. 1**), two subnuclei heavily involved in top-down regulation of fear and extinction(26,27,29). The mPFC is primarily (80-90%) glutamatergic(26); however, immunohistochemistry revealed that only 5-15% of AT2R-expressing cells in the mPFC are excitatory (**Fig. 2**), indicating that their function is likely through modulating the action of interneurons. Though the smaller (10-20%) class of mPFC neurons, several overlapping and non-overlapping interneuron types are vital to modulating the activity of excitatory projection neurons to regulate fear memory and extinction(26,36,38,39,41,43,61). However, these populations often have opposing or additive effects on mPFC output. For example, the two main interneuron populations, parvalbumin and somatostatin, are both necessary for fear memory processing(36,39) – parvalbumin interneurons, however, target the cell body of excitatory projecting cells to control neuronal output, while somatostatin interneurons target the apical dendrite to control inputs onto projecting cells(62). There is also a large degree of heterogeneity within interneuron subtypes, as illustrated by somatostatin interneurons which can be broadly classified into Martinotti cells, the primary subtype in L2/3, L5, and L6 that preferentially synapse onto excitatory projecting cells, or non-Martinotti cells, which are primarily in L4 and synapse onto parvalbumin interneurons(42,62–64). It is therefore essential to characterize which mPFC interneuron subtypes AT2R is expressed on to understand its mechanism of action.

Though AT2R has been studied on both interneuron(13) and non-interneuron(11,55) populations, its functional role and expression in the mPFC has never been examined. After determining sparse excitatory co-expression (**Fig. 2**), we stained AT2R-cre/tdTomato brains for five interneuron markers (parvalbumin, somatostatin, nNos, calbindin, and calretinin) to elucidate its role in mPFC interneuron microcircuitry. In both males and females, AT2R primarily colocalized with somatostatin-expressing interneurons, and females had greater co-expression with somatostatin than males across the PL and IL (**Fig. 4**), indicating that AT2R preferentially modulates the activity of somatostatin interneurons, a key population in regulating cognition(38,65,66) and fear learning(39–41). The minimal levels of AT2R colocalization with other interneuron populations (**Fig. 3**) may inform the type of somatostatin cell that AT2R is found on; denser levels of AT2R/calbindin co-staining combined with lower co-expression with calretinin and nNos interneurons indicates that AT2R is primarily found on excitatory cell-projecting Martinotti cells(42,64,67,68), though this conclusion is limited by the lack of AT2R-somatostatin-interneuron triple immunostaining. Additionally, the distribution of somatostatin colocalization in females in layers 5 and 2/3 (**Fig. 4**) indicates that mPFC AT2R likely has a role in the top-down control of fear and reward processing(39,40). Though we did not assess AT2R-dependent electrophysiology in the present study, AT2R activation has been previously shown to decrease spontaneous neural activity through the facilitation of potassium channel activity(15,21,22,55,69) – we therefore proposed that AT2R activation decreases somatostatin inhibition onto excitatory projection neurons, facilitating mPFC top-down control of fear. Our IHC findings (**Fig. 4**) combined with the known sex-biased effects of brain AT2R (23,52) lead us to additionally hypothesize that AT2R would have a greater effect on fear memory in female mice.

Recent studies from our lab and others have demonstrated AT2R’s role in fear extinction(11,18,70). This, as well as its direct translation to exposure-based PTSD therapies in humans(33,44,45), showed extinction learning as an ideal model for assessing the functional effect of AT2R on fear memory. We used male and female AT2R-flox mice to investigate the effect of selective mPFC AT2R deletion on extinction learning, using three extinction tests across 7 days to examine both short- and long-term memory. Consistent with previous studies on sex-specific effects(23,24,52,71), mPFC AT2R deletion significantly delayed short- and long-term extinction learning in females with no effect on fear acquisition or extinction in males (**Fig. 5**). Interestingly, while mPFC AT2R deletion did not impact locomotion or approach/avoidance behavior in males or females, it increased exploratory behavior in females (**Fig. 6**). These effects of AT2R deletion on extinction learning and exploration support our proposed AT2R-somatostatin functional mechanism, as mPFC somatostatin interneurons have been shown to encode fear memory(39–41) and regulate stress in a sex-dependent manner(43). Further, increased somatostatin signaling, which we propose occurs with AT2R deletion, increases exploratory behavior in mice(65).

The behavioral effects that we see with mPFC AT2R deletion may additionally be dependent on a TrkB/BDNF signaling pathway, as there is emerging evidence that cortical AT2R is dependent on co-expression with TrkB to influence learned fear(18,70,72). Further studies will directly interrogate the electrophysiological impact of AT2R activation or deletion on somatostatin interneuron activity and mPFC projections, as well as whether chemogenetic activation of AT2R-expressing mPFC interneurons can rescue or facilitate fear extinction.

In summary, these results identify an AT2R-expressing somatostatin interneuron population throughout the mPFC, with denser levels of co-expression in females than in males. Additionally, we show that AT2R deletion from the mPFC, while ineffective in males, prevents short-term and delays long-term extinction learning in females. Taken together, these results provide evidence for a novel sex-specific mPFC circuit where activation of AT2R on somatostatin interneurons decreases inhibition onto excitatory projection neurons, thereby increasing top-down regulation of downstream fear- and freezing-associated regions such as the amygdala, the thalamus, and the brainstem (**Fig. 7**). These results offer valuable insight into the neurobiological mechanisms of brain AT2R and top-down mPFC control of conditioned fear. They additionally increase the understanding of circuitry and function of the brain RAS in disordered fear learning, which may lead to improved therapeutics for the treatment of PTSD.

**Figure 7.**
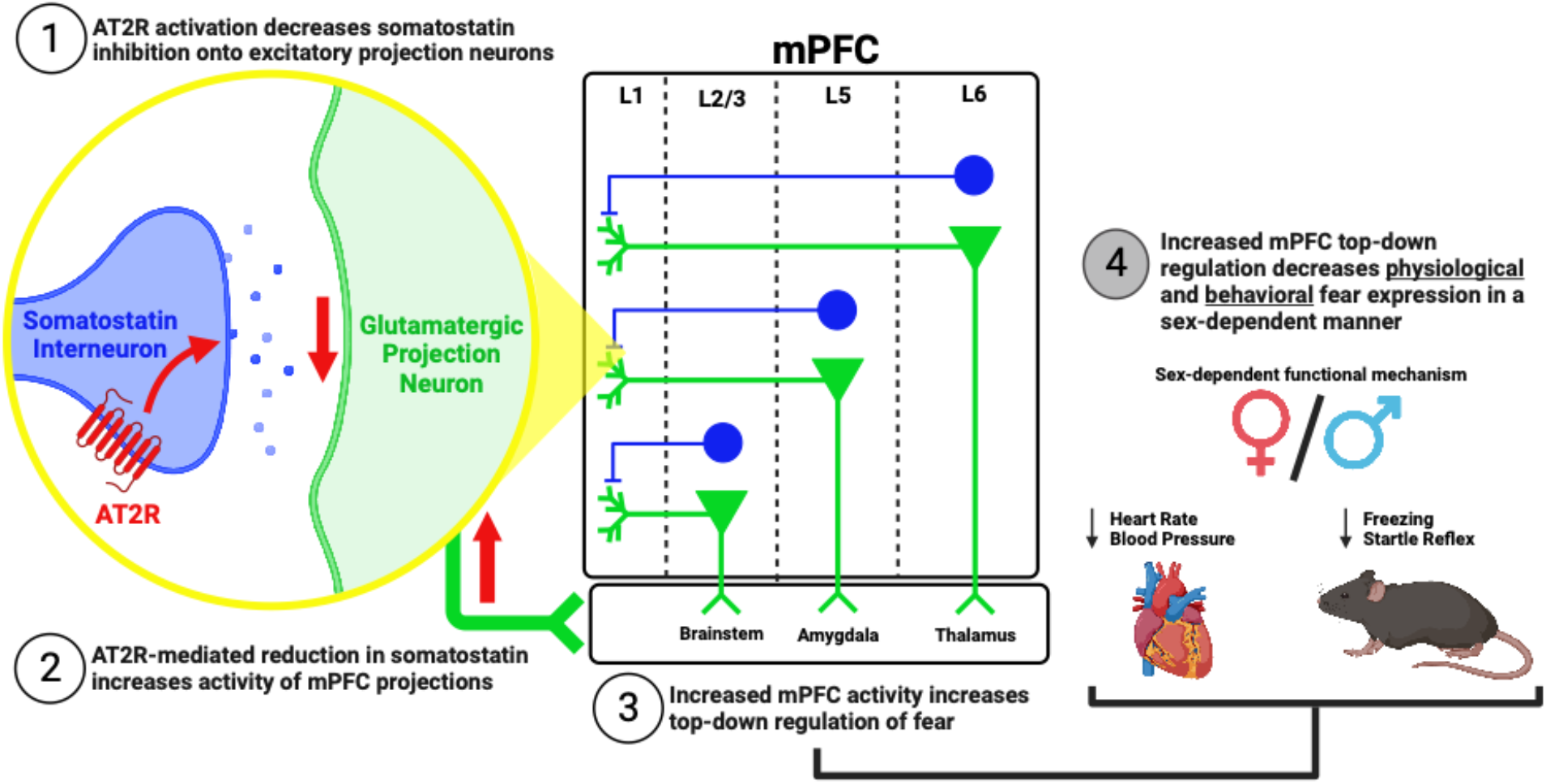
Proposed circuit for mPFC AT2R+ mechanism. **1.** Activation of AT2R on somatostatin interneurons in the PL and IL of the mPFC decreases the activity of these interneurons. **2.** This decreases inhibition of glutamatergic projection neurons, increasing their firing to downstream regions. **3.** Because of preferential somatostatin inhibition on brainstem (L2/3), amygdala (L5), and thalamic (L6) projecting excitatory neurons, mPFC top-down regulation is increased to these regions. **4.** We hypothesize that activation of mPFC AT2Rs, via increased top-down regulation, will decrease behavioral (freezing, startle) and cardioautonomic (heart rate, blood pressure) measures of fear in a sex-dependent manner.

## Acknowledgements

The authors would like to thank Drs. Eric G. Krause and Annette D. de Kloet for generously providing the AT2R-cre and AT2R-flox mouse lines. Furthermore, we would like to thank and acknowledge Drs. Cheryl Clarkson and Anastas Popratiloff and the GW Nanofabrication and Imaging Center. Finally, we thank the animal care and veterinary staff at the George Washington University for maintaining the health and well-being of our research subjects. Figures were created with BioRender.com.

## Funding

This research and PJM was supported by NIH1R01HL137103-01A1 and Congressionally Directed Medical Research Programs (CDMRP) PR210574

## Conflicts of interest statement

The authors declare no conflict of interest.

## Notes

### Competing Interest Statement

The authors have declared no competing interest.

## REFERENCES

1. Lancaster C, Teeters J, Gros D, Back S (2016): Posttraumatic Stress Disorder: Overview of Evidence-Based Assessment and Treatment. J Clin Med 5: 105.

2. Edmondson D, von Känel R (2017): Post-traumatic stress disorder and cardiovascular disease. Lancet Psychiatry Vl: 320–329.

3. Brudey C, Park J, Wiaderkiewicz J, Kobayashi I, Mellman TA, Marvar PJ (2015): Autonomic and inflammatory consequences of posttraumatic stress disorder and the link to cardiovascular disease. Am J Physiol Regul Integr Comp Physiol 309: R315–321.

4. Kimerling R, Allen MC, Duncan LE (2018): Chromosomes to Social Contexts: Sex and Gender Differences in PTSD. Curr Psychiatry Rep 20: 114.

5. Seligowski AV, Duffy LA, Merker JB, Michopoulos V, Gillespie CF, Marvar PJ, et al. (2021): The renin-angiotensin system in PTSD: a replication and extension. Neuropsychopharmacol Off Publ Am Coll Neuropsychopharmacol 46: 750–755.

6. Khoury NM, Marvar PJ, Gillespie CF, Wingo A, Schwartz A, Bradley B, et al. (2012): The Renin-Angiotensin Pathway in Posttraumatic Stress Disorder. J Clin Psychiatry 73: 849– 855.

7. Marvar PJ, Goodman J, Fuchs S, Choi DC, Banerjee S, Ressler KJ (2014): Angiotensin type 1 receptor inhibition enhances the extinction of fear memory. Biol Psychiatry 75: 864–872.

8. Swiercz A, Iyer L, Yu Z, Edwards A, Prashant NM, Nguyen BN, et al. (2020): Evaluation of an Angiotensin Type 1 Receptor Blocker on the Reconsolidation of Fear Memory. Biol Psychiatry 87: S171–S172.

9. Swiercz AP, Seligowski AV, Park J, Marvar PJ (2018): Extinction of Fear Memory Attenuates Conditioned Cardiovascular Fear Reactivity. Front Behav Neurosci 12: 276.

10. Jackson L, Eldahshan W, Fagan SC, Ergul A (2018): Within the Brain: The Renin Angiotensin System. Int J Mol Sci 19. 10.3390/ijms19030876

11. Yu Z, Swiercz AP, Moshfegh CM, Hopkins L, Wiaderkiewicz J, Speth RC, et al. (2019): Angiotensin II Type 2 Receptor–Expressing Neurons in the Central Amygdala Influence Fear-Related Behavior. Biol Psychiatry 86: 899–909.

12 de Kloet AD, Wang L, Ludin JA, Smith JA, Pioquinto DJ, Hiller H, et al. (2016): Reporter mouse strain provides a novel look at angiotensin type-2 receptor distribution in the central nervous system. Brain Struct Funct 221: 891–912.

13 de Kloet AD, Pitra S, Wang L, Hiller H, Pioquinto DJ, Smith JA, et al. (2016): Angiotensin Type-2 Receptors Influence the Activity of Vasopressin Neurons in the Paraventricular Nucleus of the Hypothalamus in Male Mice. Endocrinology 157: 3167–3180.

14. Mohammed M, Sumners C, Sheng W, Harden SW, Frazier CJ, Johnson D, et al. (2020): Angiotensin AT2 receptors in the solitary tract nucleus lower blood pressure via inhibition of GABA signaling. FASEB J 34: 1–1.

15. Sumners C, Alleyne A, Rodríguez V, Pioquinto DJ, Ludin JA, Kar S, et al. (2020): Brain angiotensin type-1 and type-2 receptors: cellular locations under normal and hypertensive conditions. Hypertens Res 43: 281–295.

16. de Kloet AD, Steckelings UM, Sumners C (2017): Protective Angiotensin Type 2 Receptors in the Brain and Hypertension. Curr Hypertens Rep 19. 10.1007/s11906-017-0746-x

17. Faria-Costa G, Leite-Moreira A, Henriques-Coelho T (2014): Cardiovascular effects of the angiotensin type 2 receptor. Rev Port Cardiol Engl Ed 33: 439–449.

18. Diniz CRAF, Casarotto PC, Fred SM, Biojone C, Castrén E, Joca SRL (2018): Antidepressant-like effect of losartan involves TRKB transactivation from angiotensin receptor type 2 (AGTR2) and recruitment of FYN. Neuropharmacology 135: 163–171.

19. Okuyama S, Sakagawa T, Chaki S, Imagawa Y, Ichiki T, Inagami T (1999): Anxiety-like behavior in mice lacking the angiotensin II type-2 receptor. Brain Res 821: 150–159.

20. Kerr DS, Bevilaqua LRM, Bonini JS, Rossato JI, Köhler CA, Medina JH, et al. (2004): Angiotensin II blocks memory consolidation through an AT2 receptor-dependent mechanism. Psychopharmacology (Berl) 179: 529–535.

21. Gao L, Zucker IH (2011): AT2 receptor signaling and sympathetic regulation. Curr Opin Pharmacol 11: 124–130.

22. Xiong H, Marshall KC (1990): Angiotensin II modulation of glutamate excitation of locus coeruleus neurons. Neurosci Lett 118: 261–264.

23. Barsha G, Walton SL, Kwok E, Denton KM (2019): Sex Differences in the Role of the Angiotensin Type 2 Receptor in the Regulation of Blood Pressure. Sex Differences in Cardiovascular Physiology and Pathophysiology. Elsevier, pp 73–103.

24. Armando I, Jezova M, Juorio AV, Terrón JA, Falcón-Neri A, Semino-Mora C, et al. (2002): Estrogen upregulates renal angiotensin II AT _2_ receptors. Am J Physiol-Ren Physiol 283: F934–F943.

25. Sakata A, Mogi M, Iwanami J, Tsukuda K, Min L-J, Fujita T, et al. (2009): Sex-different effect of angiotensin II type 2 receptor on ischemic brain injury and cognitive function. Brain Res 1300: 14–23.

26. Anastasiades PG, Carter AG (2021): Circuit organization of the rodent medial prefrontal cortex. Trends Neurosci 44: 550–563.

27. Arnsten AFT (2015): Stress weakens prefrontal networks: molecular insults to higher cognition. Nat Neurosci 18: 1376–1385.

28. Arnsten AFT, Raskind MA, Taylor FB, Connor DF (2015): The effects of stress exposure on prefrontal cortex: Translating basic research into successful treatments for post-traumatic stress disorder. Neurobiol Stress 1: 89–99.

29. Courtin J, Bienvenu TCM, Einarsson EÖ, Herry C (2013): Medial prefrontal cortex neuronal circuits in fear behavior. Neuroscience 240: 219–242.

30. Gilmartin MR, Balderston NL, Helmstetter FJ (2014): Prefrontal cortical regulation of fear learning. Trends Neurosci 37: 455–464.

31. Giustino TF, Maren S (2015): The Role of the Medial Prefrontal Cortex in the Conditioning and Extinction of Fear. Front Behav Neurosci 9. 10.3389/fnbeh.2015.00298

32. Fucich EA, Paredes D, Saunders MO, Morilak DA (2018): Activity in the Ventral Medial Prefrontal Cortex Is Necessary for the Therapeutic Effects of Extinction in Rats. J Neurosci 38: 1408–1417.

33. Zhou F, Geng Y, Xin F, Li J, Feng P, Liu C, et al. (2019): Human Extinction Learning Is Accelerated by an Angiotensin Antagonist via Ventromedial Prefrontal Cortex and Its Connections With Basolateral Amygdala. Biol Psychiatry 86: 910–920.

34. Hiscox LV, Sharp TH, Olff M, Seedat S, Halligan SL (2023): Sex-Based Contributors to and Consequences of Post-traumatic Stress Disorder. Curr Psychiatry Rep 25: 233–245.

35. Pooley AE, Benjamin RC, Sreedhar S, Eagle AL, Robison AJ, Mazei-Robison MS, et al. (2018): Sex differences in the traumatic stress response: PTSD symptoms in women recapitulated in female rats. Biol Sex Differ 9: 31.

36. Binette AN, Liu J, Bayer H, Crayton KL, Melissari L, Sweck SO, Maren S (2023): Parvalbumin-positive interneurons in the medial prefrontal cortex regulate stress-induced fear extinction impairments in male and female rats. J Neurosci JN-RM-1442–22.

37. Chung DW, Wills ZP, Fish KN, Lewis DA (2017): Developmental pruning of excitatory synaptic inputs to parvalbumin interneurons in monkey prefrontal cortex. Proc Natl Acad Sci 114: E629–E637.

38. Yang S-S, Mack NR, Shu Y, Gao W-J (2021): Prefrontal GABAergic Interneurons Gate Long-Range Afferents to Regulate Prefrontal Cortex-Associated Complex Behaviors. Front Neural Circuits 15: 716408.

39. Cummings KA, Clem RL (2020): Prefrontal somatostatin interneurons encode fear memory. Nat Neurosci 23: 61–74.

40. Cummings KA, Bayshtok S, Dong TN, Kenny PJ, Clem RL (2022): Control of fear by discrete prefrontal GABAergic populations encoding valence-specific information. Neuron 110: 3036–3052.e5.

41 Koppensteiner P, Von Itter R, Melani R, Galvin C, Lee FS, Ninan I (2019): Diminished Fear Extinction in Adolescents Is Associated With an Altered Somatostatin Interneuron– Mediated Inhibition in the Infralimbic Cortex. Biol Psychiatry 86: 682–692.

42. Riedemann T (2019): Diversity and Function of Somatostatin-Expressing Interneurons in the Cerebral Cortex. Int J Mol Sci 20: 2952.

43. Girgenti MJ, Wohleb ES, Mehta S, Ghosal S, Fogaca MV, Duman RS (2019): Prefrontal cortex interneurons display dynamic sex-specific stress-induced transcriptomes. Transl Psychiatry 9: 292.

44. Andero R, Ressler KJ (2012): Fear extinction and BDNF: translating animal models of PTSD to the clinic. Genes Brain Behav 11: 503–512.

45. Cain CK (2023): Beyond Fear, Extinction, and Freezing: Strategies for Improving the Translational Value of Animal Conditioning Research. Berlin, Heidelberg: Springer Berlin Heidelberg. 10.1007/7854_2023_434

46. Shansky RM (2015): Sex differences in PTSD resilience and susceptibility: Challenges for animal models of fear learning. Neurobiol Stress 1: 60–65.

47. Fenster RJ, Lebois LAM, Ressler KJ, Suh J (2018): Brain circuit dysfunction in post-traumatic stress disorder: from mouse to man. Nat Rev Neurosci 19: 535–551.

48. Dulka BN, Bagatelas ED, Bress KS, Grizzell JA, Cannon MK, Whitten CJ, Cooper MA (2020): Chemogenetic activation of an infralimbic cortex to basolateral amygdala projection promotes resistance to acute social defeat stress. Sci Rep 10: 6884.

49. Singewald N, Schmuckermair C, Whittle N, Holmes A, Ressler KJ (2015): Pharmacology of cognitive enhancers for exposure-based therapy of fear, anxiety and trauma-related disorders. Pharmacol Ther 149: 150–190.

50. Day HLL, Stevenson CW (2020): The neurobiological basis of sex differences in learned fear and its inhibition. Eur J Neurosci 52: 2466–2486.

51. Lazaroni TLDN, Bastos CP, Moraes MFD, Santos RS, Pereira GS (2016): Angiotensin-(1-7)/Mas axis modulates fear memory and extinction in mice. Neurobiol Learn Mem 127: 27–33.

52. Sakata A, Mogi M, Iwanami J, Tsukuda K, Min L-J, Fujita T, et al. (2009): Sex-different effect of angiotensin II type 2 receptor on ischemic brain injury and cognitive function. Brain Res 1300: 14–23.

53. Jing F, Mogi M, Sakata A, Iwanami J, Tsukuda K, Ohshima K, et al. (2012): Direct Stimulation of Angiotensin II Type 2 Receptor Enhances Spatial Memory. J Cereb Blood Flow Metab 32: 248–255.

54. Okuyama S, Sakagawa T, Chaki S, Imagawa Y, Ichiki T, Inagami T (1999): Anxiety-like behavior in mice lacking the angiotensin II type-2 receptor. Brain Res 821: 150–159.

55. Mohammed M, Johnson DN, Wang LA, Harden SW, Sheng W, Spector EA, et al. (2022): Targeting angiotensin type-2 receptors located on pressor neurons in the nucleus of the solitary tract to relieve hypertension in mice. Cardiovasc Res 118: 883–896.

56. Yu Z, Kisner A, Bhatt A, Polter AM, Marvar PJ (2023): Central amygdala angiotensin type 1 receptor (Agtr1) expressing neurons contribute to fear extinction. Neuropharmacology 229: 109460.

57. Stout DM, Risbrough VB (2019): Angiotensin II Signaling and Fear Extinction: Translational Evidence and Novel Receptor Targets. Biol Psychiatry 86: 874–876.

58. Stein MB, Jain S, Simon NM, West JC, Marvar PJ, Bui E, et al. (2021): Randomized, Placebo-Controlled Trial of the Angiotensin Receptor Antagonist Losartan for Posttraumatic Stress Disorder. Biol Psychiatry 90: 473–481.

59. Wang D, Hu S, Zhu J, Yuan J, Wu J, Zhou A, et al. (2013): Angiotensin II type 2 receptor correlates with therapeutic effects of losartan in rats with adjuvant-induced arthritis. J Cell Mol Med 17: 1577–1587.

60. He D-H, Lin J-X, Zhang L-M, Xu C-S, Xie Q (2017): Early treatment with losartan effectively ameliorates hypertension and improves vascular remodeling and function in a prehypertensive rat model. Life Sci 173: 20–27.

61. Woodward EM, Coutellier L (2021): Age- and sex-specific effects of stress on parvalbumin interneurons in preclinical models: Relevance to sex differences in clinical neuropsychiatric and neurodevelopmental disorders. Neurosci Biobehav Rev 131: 1228– 1242.

62. Shepherd GM, Grillner S (Eds.) (2018): Handbook of Brain Microcircuits, Second edition. New York, NY: Oxford University Press.

63. Liguz-Lecznar M, Dobrzanski G, Kossut M (2022): Somatostatin and Somatostatin-Containing Interneurons—From Plasticity to Pathology. Biomolecules 12: 312.

64. Nigro MJ, Hashikawa-Yamasaki Y, Rudy B (2018): Diversity and Connectivity of Layer 5 Somatostatin-Expressing Interneurons in the Mouse Barrel Cortex. J Neurosci 38: 1622– 1633.

65. Brockway DF, Griffith KR, Aloimonos CM, Clarity TT, Moyer JB, Smith GC, et al. (2023): Somatostatin peptide signaling dampens cortical circuits and promotes exploratory behavior. Cell Rep 42: 112976.

66. Kupferschmidt DA, Cummings KA, Joffe ME, MacAskill A, Malik R, Sánchez-Bellot C, et al. (2022): Prefrontal Interneurons: Populations, Pathways, and Plasticity Supporting Typical and Disordered Cognition in Rodent Models. J Neurosci 42: 8468–8476.

67. Yavorska I, Wehr M (2016): Somatostatin-Expressing Inhibitory Interneurons in Cortical Circuits. Front Neural Circuits 10. 10.3389/fncir.2016.00076

68. McKlveen JM, Moloney RD, Scheimann JR, Myers B, Herman JP (2019): “Braking” the Prefrontal Cortex: The Role of Glucocorticoids and Interneurons in Stress Adaptation and Pathology. Biol Psychiatry 86: 669–681.

69. Steckelings UM, Kloet A de, Sumners C (2017): Centrally Mediated Cardiovascular Actions of the Angiotensin II Type 2 Receptor. Trends Endocrinol Metab 28: 684–693.

70. Laukkanen L, Diniz CRAF, Foulquier S, Prickaerts J, Castrén E, Casarotto PC (2021): Facilitation of TRKB Activation by the Angiotensin II Receptor Type-2 (AT2R) Agonist C21. Pharmaceuticals 14: 773.

71. Sullivan JC (2008): Sex and the renin-angiotensin system: inequality between the sexes in response to RAS stimulation and inhibition. Am J Physiol-Regul Integr Comp Physiol 294: R1220–R1226.

72. Goel R, Bhat SA, Hanif K, Nath C, Shukla R (2018): Angiotensin II Receptor Blockers Attenuate Lipopolysaccharide-Induced Memory Impairment by Modulation of NF-κB-Mediated BDNF/CREB Expression and Apoptosis in Spontaneously Hypertensive Rats. Mol Neurobiol 55: 1725–1739.

